# The fitness effects of spontaneous mutations nearly unseen by selection in a bacterium with multiple chromosomes

**DOI:** 10.1101/060483

**Authors:** Marcus M. Dillon, Vaughn S. Cooper

## Abstract

Mutation accumulation (MA) experiments employ the strategy of minimizing the population size of evolving lineages to greatly reduce effects of selection on newly arising mutations. Thus, most mutations fix within MA lines independently of their fitness effects. This approach, more recently combined with genome sequencing, has detailed the rates, spectra, and biases of different mutational processes. However, a quantitative understanding of the fitness effects of mutations virtually unseen by selection has remained an untapped opportunity. Here, we analyzed the fitness of 43 sequenced MA lines of the multi-chromosome bacterium *Burkholderia cenocepacia* that had each undergone 5554 generations of MA and accumulated an average of 6.73 spontaneous mutations. Most lineages exhibited either neutral or deleterious fitness in three different environments in comparison with their common ancestor. The only mutational class that was significantly overrepresented in lineages with reduced fitness was the loss of the plasmid, though nonsense mutations, missense mutations, and coding insertion-deletion mutations were also overrepresented in MA lineages whose fitness had significantly declined. Although the overall distribution of fitness effects was similar between the three environments, the magnitude and even the sign of the fitness of a number of lineages changed with the environment, demonstrating that the fitness of some genotypes was environmentally dependent. These results present an unprecedented picture of the fitness effects of spontaneous mutations in a bacterium with multiple chromosomes and provide greater quantitative support of the theory that the vast majority of spontaneous mutations are neutral or deleterious.

## INTRODUCTION

The extent to which spontaneous mutations contribute to evolutionary change largely depends on their rates and fitness effects. Both parameters are fundamental to several evolutionary problems, including the preservation of genetic variation (Charlesworth *et al.* 1993, 2009; Charlesworth and Charlesworth 1998), the evolution of recombination (Muller 1964; Kondrashov 1988; Otto and Lenormand 2002; Roze and Blanckaert 2014), the evolution of mutator alleles (Sniegowski *et al.* 1997; Tenaillon *et al.* 1999), and deleterious mutation accumulation in small populations (Lande 1994; Lynch *et al.* 1995, 1999; Schwander and Crespi 2009). Many studies have now obtained direct and robust estimates of mutation rates and spectra across diverse organisms, but our understanding of the fitness effects of spontaneous mutations remains limited to mostly indirect estimates in classic model organisms (Eyre-Walker and Keightley 2007).

Mutation accumulation (MA) experiments provide the opportunity to quantify properties of the fitness of spontaneous mutations that have not been exposed to the sieve of natural selection. Specifically, MA experiments limit the efficiency of natural selection by passaging replicate lineages through repeated single cell bottlenecks. These lineages accumulate mutations independently over several thousand generations, and the magnitude and variance in fitness between lineages can be used to estimate several properties of the distribution of fitness effects (Halligan and Keightley 2009). MA studies have been used to characterize the fitness effects of spontaneous mutations in *Drosophila melanogaster* (Bateman 1959; Mukai 1964; Keightley 1994; Fry *et al.* 1999), *Arabidopsis thaliana* (Schultz *et al.* 1999; Shaw *et al.* 2000, 2002), *Caenorhabditis elegans* (Keightley and Caballero 1997; Vassilieva *et al.* 2000; Estes *et al.* 2004; Katju *et al.* 2015), *Saccharomyces cerevisiae* (Wloch *et al.* 2001; Zeyl and de Visser 2001; Dickinson 2008; Jasmin and Lenormand 2015), *Escherichia coli* (Kibota and Lynch 1996; Trindade *et al.* 2010), and other microbes (Heilbron *et al.* 2014; Kraemer *et al.* 2015). Results from these studies have occasionally been inconsistent, but the majority of results suggest that most spontaneous mutations have mild effects (Eyre-Walker and Keightley 2007; Halligan and Keightley 2009; Agrawal and Whitlock 2012; Heilbron *et al.* 2014), that deleterious mutations far outnumber beneficial mutations (Keightley and Lynch 2003; Silander *et al.* 2007; Eyre-Walker and Keightley 2007), and that the distribution of effects of deleterious mutations is complex and multimodal (Zeyl and de Visser 2001; Eyre-Walker and Keightley 2007).

A more powerful approach for studying the fitness effects of spontaneous mutations is to pair MA experiments with whole-genome sequencing (MA-WGS), so that both the genetic basis and fitness effects of a collection of mutations can be known. MA-WGS studies have been conducted in a diverse array of bacteria, generating a growing database of naturally accumulated mutations that has dramatically improved estimates of mutation rates and spectra (Lee *et al.* 2012; Sung *et al.* 2012, 2015; Heilbron *et al.* 2014; Long *et al.* 2014, 2015; Dillon *et al.* 2015; Foster *et al.* 2015; Dettman *et al.* 2016). Yet, only one of these studies has also characterized the fitness of MA-WGS lines (Heilbron *et al.* 2014), and this study was conducted with mutator lineages, which have altered base-substitution and indel biases (Lee *et al.* 2012; Sung *et al.* 2015), and produce hundreds of mutations per line. Our understanding of the fitness effects of spontaneous mutations would benefit greatly from more direct estimates of fitness derived from MA lineages that harbor fewer known mutations.

Here, we measured the relative fitness of 43 fully sequenced MA lineages derived from *B. cenocepacia* HI2424 in three laboratory environments after they had been evolved in the near absence of natural selection for 5554 generations. Following the MA experiment, each lineage harbored a total mutational load of 2 to 14 spontaneous mutations, including base-substitution mutations (bpsms), insertion-deletion mutations (indels), and whole-plasmid deletions. By correlating the relative fitness of these MA lineages with the particular mutations that they harbor, we present a detailed picture of the fitness effects of spontaneous mutations, and precise estimates of deleterious mutation rates and fitness effects in a clinically important gamma-proteobacterium with multiple chromosomes.

## MATERIALS AND METHODS

**Bacterial strains and culture conditions.** All mutation accumulation experiments were founded from a single colony of *Burkholderia cenocepacia* HI2424, which was isolated from the soil and only passaged in the laboratory during isolation (Coenye and LiPuma 2003). As a member of the diverse *B. cepacia* complex, *Burkholderia cenocepacia* can form highly resistant biofilms and has been associated with persistent lung infections in patients with cystic fibrosis (Mahenthiralingam *et al.* 2005; Traverse *et al.* 2013). The genome of *B. cenocepacia* HI2424 has been fully sequenced and is composed of three chromosomes (Chr1: 3.48-Mb, 3253 genes; Chr2: 3.00-Mb, 2709 genes; Chr3: 1.06Mb, 929 genes) and a plasmid (0.164-Mb, 159 genes), though the third chromosome can be eliminated under some conditions in some strains (Agnoli *et al.* 2012). To facilitate relative fitness assays, we competed all *B. cenocepacia* HI2424 strains derived from the MA experiment with a *B. cenocepacia* HI2424 Lac+ strain, which is isogenic to *B. cenocepacia* HI2424, except for the introduction of the *lacZ* gene at the *attTn7* site, which causes colonies to turn blue when exposed to 5-bromo-4-chloro-indolyl-β-galactopyranoside (X-gal) (Choi *et al.* 2005).

MA experiments were conducted on tryptic soy agar plates (TSA) (30 g/liter tryptic soy broth powder, 15 g/liter agar) and were incubated at 37°. At the conclusion of the MA experiment, frozen stocks were prepared by growing a single colony from each lineage overnight in 5ml of tryptic soy broth (TSOY) (30 g/liter tryptic soy broth powder) at 37° and freezing at −80° in 8% DMSO. All relative fitness assays were conducted in 18 × 150mm glass capped tubes with 5ml of liquid medium and were maintained at 37° in a roller drum (30 rpm). Relative fitness of each lineage was assayed in three different environments. First, we conducted relative fitness assays in TSOY, a medium that mimics the conditions of the MA experiment and is expected to be very permissive. Second, we conducted relative fitness assays in M9 Minimal Medium supplemented with 0.3% casamino acids (M9MM+CAA) (3 g/liter casamino acid powder, 1 g/liter glucose, 6 g/liter sodium phosphate dibasic anhydrous, 3 g/liter potassium phosphate monobasic, 1 g/liter ammonium chloride, 0.5 g/liter sodium chloride, 0.1204 g/liter magnesium sulfate, 0.0147 g/liter calcium chloride), a medium that is more nutrient restrictive than TSOY, but contains all essential amino acids except tryptophan. Lastly, we conducted relative fitness assays in M9 Minimal Medium (M9MM) (1 g/liter glucose, 6 g/liter sodium phosphate dibasic anhydrous, 3 g/liter potassium phosphate monobasic, 1 g/liter ammonium chloride, 0.5 g/liter sodium chloride, 0.1204 g/liter magnesium sulfate, 0.0147 g/liter calcium chloride), which is a fully defined medium that is more restrictive than either TSOY or M9MM+CAA. Serial passaging during fitness assays was performed using 100-fold dilutions, so all relative fitness assays were conducted over the same number of generations, despite the moderate different carrying capacities in these mediums. All dilutions were performed using phosphate buffer saline (PBS) (80 g/liter NaCl, 2 g/liter KCl, 14.4 g/liter Na2HPO4 • 2H2O, 2.4 g//liter KH_2_PO_4_) in 96-well plates.

**MA-WGS process.** The mutation accumulation experiment that generated the mutational load for this study has been reported previously (Dillon *et al.* 2015). Briefly, seventy-five independent lineages were founded from a single colony of *B. cenocepacia* HI2424 and independently propagated every 24 hours onto a fresh TSA plate for 217 days. Daily generations were estimated monthly by taking a single representative colony from each lineage following 24 hours of growth, placing it in 2 ml of PBS, then serially diluting and spread plating it on TSA. The number of viable cells in each colony was then used to calculate the number of generations elapsed between each transfer, and the average number of generations across all lineages was used as the number of generations per day for that entire month. By multiplying the number of generations per day for each month by the number of days in that month, then summing these totals over the course of the whole experiment, we calculated the total number of generations elapsed per MA lineage over the course of the MA-experiment.

As we have described previously, genomic DNA was extracted from 1 ml of overnight TSB culture founded by 47 of the *B. cenocepacia* isolates that were stored at the conclusion of the MA experiment (Dillon *et al.* 2015). We used the Wizard Genomic DNA Purification kit for DNA extraction (Promega), all libraries were prepared using a modified Illumina Nextera protocol (Baym *et al.* 2015), and sequencing was performed with the 151-bp paired end platform on the Illumina HiSeq at the Hubbard Center for Genomic Studies at the University of New Hampshire. Following fastQC analysis, all reads were mapped to the *B. cenocepacia* HI2424 reference genome (LiPuma *et al.* 2002) with both the Burrows-Wheeler aligner (BWA) (Li and Durbin 2009) and Novoalign (www.novocraft.com). The average depth of coverage across all 47 lines was 43×, but the average depth of coverage in the 43 lines used in this study was 46×.

**Spontaneous mutation identification.** All bpsms were identified as described previously (Dillon *et al.* 2015). Briefly, after using a combination of SAMtools and in house perl scripts to produce all read alignments for each position in each line (Li *et al.* 2009), a three step process was used to detect putative bpsms. First, pooled reads across all lines were used to generate an ancestral consensus base at each site in the reference genome, allowing us to correct differences between the published reference genome and the ancestral colony of our MA experiment. Second, reads from the individual lines were used to generate a lineage specific consensus base at each site in the reference genome for each lineage, as long as the site was covered by at least two forward and two reverse reads, and at least 80% of the reads identified the same base. Sites that did not meet these criteria were not analyzed in the respective lineage. Third, lineage specific consensus bases for each lineage were compared to the ancestral consensus base at each site, and a putative bpsm was identified if they differed. This three-step process was carried out independently using both the BWA and Novoalign alignments, and putative bpsms were considered genuine only if both pipelines independently identified the bpsm. Despite these lenient criteria, all of the bpsms that were identified at analyzed sites in this study had considerably greater coverage and consensus than the minimal criteria (see File S1), demonstrating that these bpsms are not merely false positives in low coverage regions. The frequency of sites that were not analyzed in each lineage varied from 0.005 to 0.177. Putative bpsms in these regions were estimated by multiplying the number of unanalyzed sites in each lineage by the overall bpsm rate calculated in this study (1.31 (0.08) • 10^−10^ /bp/generation) and the number of generations of mutation accumulation in each lineage (5554) (see Table S1).

Indels and large structural variants are inherently more difficult to identify than bpsms because gaps and simple sequence repeats (SSRs) reduce the accuracy of short-read alignment algorithms. To overcome these issues, we extracted all putative indels if at least 30% of the reads that covered the site identified the exact same indel (size and motif), and the site was covered by at least two forward and two reverse reads (Dillon *et al.* 2015). These putative indels were then subject to a series of more strenuous filters based on the complexity of the region in which they were identified and the consensus between the BWA and Novoalign alignments (see File S1). Specifically, all putative indels where more than 80% of the reads identified the exact same indel in both the BWA and Novoalign alignments were considered genuine indels. For putative indels where only 30-80% of the reads identified the exact same indel, we parsed out only reads that had bases covering both the upstream and downstream region of the indel (if it was not in an SSR), and both the upstream and downstream region of the SSR (if it was in an SSR). Using this subset of reads, we reassessed the frequency of reads that identified the exact same indel, allowing us to more accurately identify indels involving the gain or loss of a single repeat within a SSR. These indels were considered genuine if more than 80% of the parsed reads identified the exact same indel and were discarded if they did not. Putative indels and other structural variants were also extracted using PINDEL, which uses paired-end information to identify insertions, deletions, inversions, tandem duplications and other structural variants (Ye *et al.* 2009). Here, indels were considered genuine if they were covered by at least six forward and six reverse reads, and at least 80% of the reads identified the exact same indel. Lastly, we analyzed the distribution of coverage between chromosomes and the 0.164-Mb plasmid to detect any chromosomal copy number variants. As with bpsms, putative indels in regions that were not analyzed were estimated by multiplying the number of unanalyzed sites in each lineage by the overall indel rate calculated in this study (2.39 11 (0.34) • 10^−11^ /bp/generation) and the number of generations of mutation accumulation in each lineage (5554) (see Table S1).

**Quantifying relative fitness.** To quantify the selection coefficients of each of the 43 derived MA lineages, we conducted three-day competitions (over ≈ 19.93 generations) between each MA lineage and our *B. cenocepacia* HI2424 Lac+ strain. These competitions were carried out independently in TSOY, M9MM+CAA, and M9MM, with four replicates being conducted for each lineage in each environment. Our MA ancestral*B. cenocepacia* HI2424 strain was also competed against *B. cenocepacia* HI2424 Lac+ as a control, with four replicates for each environment. Selection coefficients were estimated as described previously, using the relative growth of the focal MA lineage and the *B. cenocepacia* HI2424 Lac+ reference strain, normalized by the number of generations elapsed by the reference strain *(G)* (Chevin 2011; Perfeito *et al.* 2014). First, the difference in growth (Δ*r_ab_*) between the two strains was estimated as:

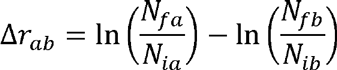

where *N_ia_* and *N_ib_* were the initial numbers of test and reference bacteria, respectively, and *N_fa_* and *N_fb_* were the final numbers of test and reference bacteria, respectively. Selection coefficients (*S_ab_*)were then calculated as:

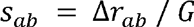

where *G,* generations elapsed by the reference strain, is equal to:

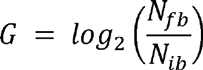

For each replicate, all 43 derived MA lineages, *B. cenocepacia* HI2424, and *B. cenocepacia* Lac+ were resurrected from frozen culture by inoculating them into 5 ml of TSOY broth and incubating overnight in a roller drum at 30 rpm. Depending on which environment was being assayed, each strain was then transferred to fresh TSOY, M9MM+CAA, or M9MM via a 10,000-fold dilution and acclimated for 24 hours at 37° and 30 rpm. Following acclimation, 44 competitions (43 MA lineages + control) were generated in the appropriate fresh medium at a 1:1 ratio via 100-fold dilution, and 30 μl from each was extracted to quantify the initial frequency of each competitor (*N_ia_, N_ib_*). Competitions were then incubated for 72 hours at 37° and 30 rpm, being transferred to fresh media every 24 hours via a 100-fold dilution. At the conclusion of the 72 hour competition, 30 μl of the final culture was extracted to quantify the final frequency of each competitor (*N_fa_, N_fb_*).

To measure the initial frequency of each competitor, the extracted culture was diluted in PBS and 100 μl of the diluted sample was plated on a TSA + X-Gal plate. Specifically, in the TSOY competitions the samples were diluted 30,000-fold, in the M9MM+CAA competitions the samples were diluted to 20,000-fold, and in the M9MM competitions the samples were diluted to 10-000-fold. Following a 48-hr incubation, the number of white and blue colonies were quantified and used to calculate *N_ia_* and *N_ib_*, respectively, after accounting for the dilutions. Final frequencies were measured in the same way, except that an additional 100-fold dilution was required for each competition because the cultures were at carrying capacity. In addition, to calculate *N_fa_* and *N_fb_*, we had to account for the dilutions that were conducted prior to plating and the two 100-fold dilutions that were conducted during the three-day competition. Importantly, the selection coefficient of the *B. cenocepacia* HI2424 MA ancestor was not significantly different from 0 in any of environments (TSOY: *s* = −0.0002 (0.0020), M9MM+CAA: *s* = +0.0075 (0.0030), M9MM: s = −0.0016 (0.0034) (SEM)).

**Statistical analysis.** All statistical analyses were performed in R Version 0.98.1091 using the Stats analysis package (R Development Core Team 2013). For independent two-tailed t-tests, all p-values were corrected for multiple comparisons using a Benjamini-Hochberg correction (Table S2), which ensures that our false positive rate remains below 5%, despite testing whether the selection coefficient differed significantly from 0 for 43 lineages in each environment (Benjamini and Hochberg 1995). Corrected p-values that were below a threshold of 0.05 were considered significant. Linear regressions were used to evaluate the correlation between the number of mutations in a lineage and its selection coefficient, as well as the correlation between the selection coefficients of lineages in different environments. Lastly, to test for effects of replicate, genotype, environment, and genotype^*^environment interaction on the fitness of each lineage, we performed an analysis of variance (ANOVA) on the cumulative dataset.

**Data Availability.** Illumina DNA sequences for the *B. cenocepacia* MA lines used in this study are available under the PRJNA326274 bioproject. File S1 contains all supplementary tables and figures referenced in text. File S2 contains all the information on the location and criteria used to identify each base-substitution and insertion-deletion mutation identified in this study. All strains are available upon request.

## RESULTS

We previously reported the rate and molecular spectrum of spontaneous mutations in wild-type *B. cenocepacia*, as determined from the cumulative results of a MA-WGS experiment involving 47 replicate lineages derived from *B. cenocepacia* HI2424 (Dillon *et al.* 2015). Each lineage was passaged through daily single-cell bottlenecks for 217 days, resulting in approximately 5554 generations of MA per lineage. The average number of generations of growth per day within a colony declined from 26.16 (0.06) to 24.92 (0.07) (SEM) over the course of the experiment, suggesting that some of the accumulated mutations had deleterious fitness effects. Here, we present a detailed picture of the fitness effects of the spontaneous mutations accumulated at the conclusion of this MA-WGS experiment using 43 replicate lineages, as the remaining four lineages were discarded because of a lack of sufficient coverage in the WGS data.

The properties of the mutations found in these 43 lineages are consistent with constant mutation rates and limited selection over the course of our experiment. Neither the distribution of bpsms or indels across lineages differed significantly from a Poisson distribution (bpsms: χ^2^ = 3.46, p = 0.94; indels: χ^2^ = 0.28, p = 0.96), signifying that mutation rates did not vary across the lineages. The ratio of synonymous to nonsynonymous bpsms also did not differ from the expected ratio based on the codon-usage and %GC content at synonymous and nonsynonymous sites in *B. cenocepacia* HI2424 (χ^2^ = 0.78, d.f. = 1, p = 0.38), which suggests minimal purifying selection. Further, the lack of genetic parallelism in the bpsm spectra across lineages (see File S1) is inconsistent with positive selection acting on these lines. Although both bpsms and indels were observed more frequently than expected in non-coding DNA (bpsms: χ^2^ = 2.19, d.f. = 1, p = 0.14; indels: χ^2^ = 45.816, d.f. = 1, p < 0.0001), a pattern consistent with purifying selection, this pattern could be generated by selection against coding mutations, preferential mismatch repair in coding regions, or the mutation prone nature of repetitive DNA in non-coding regions (Lee *et al.* 2012; Heilbron *et al.* 2014; Dillon *et al.* 2015). In any event, we estimate that the threshold selection coefficient below which genetic drift will overpower natural selection, as determined by *N_E_* × *s* =1 in haploid organisms, is 0.08 (Dillon *et al.* 2015). Thus, while a small class of adaptive or deleterious mutations with effects in excess of *s* = 0.08 will be subject to the biases of natural selection (Kimura 1983; Elena *et al.* 1998; Zeyl and de Visser 2001; Hall *et al.* 2008), the vast majority of mutations that were observed in this study likely fixed irrespective of their fitness effects.

**Genetic basis of spontaneous mutations.** The spontaneous bpsms and indels reported here are similar to those reported previously (Dillon *et al.* 2015), with a few exceptions. First, we allowed for bpsms to be called in more than one lineage, resulting in the addition of two bpsms. These bpsms are assumed to have occurred in the ancestral colony, but their presence in each lineage must be documented to accurately quantify the relationship between the fitness of each lineage and its mutational load. Second, we were able to confidently identify nine additional indels that occurred in simple sequence repeats by using an approach that considers only reads anchored on both sides of the repeat (see Methods; File S1). These indels may contribute substantively to the mutational load of the lineages in which they occur, so they were important to include in this study. Lastly, we did not analyze four of the lineages from the previous study because less than 80% of their genomes had sufficient coverage to be analyzed for the presence of bpsms and indels (see Methods). This low coverage would render us blind to a considerable portion of the mutational load in these lineages, which warranted their exclusion.

Among the 43 MA lineages, 233 bpsms, 42 short indels, and 4 plasmid-loss events were identified. The most common class of bpsms was missense bpsms (141), followed by synonymous bpsms (49), intergenic bpsms (37), and nonsense bpsms (6). Among indels, coding indels involving only a single gene (22) were slightly more common than intergenic indels (20), while loss of the 0.16-Mb plasmid, which encodes 157 genes, was observed in 4 lineages (see Figure 1). False-negative rates were estimated in each lineage as the number of sites that were not analyzed for mutations, multiplied by the product of experiment-wide bpsm and indel rates per base-pair per generation and the number of generations experienced by each lineage. Given that an average of 95.20 % (0.01) of each genome was sequenced to sufficient depth to analyze both bpsms and indels, we estimate that an average of only 0.25 (0.03) additional bpsms and 0.05 (0.01) additional indels would have been identified per lineage if the entire genome was analyzed (SEM) (see Table S1). Overall, mutations were not uniformly distributed across the 43 MA lineages, allowing us to analyze which mutation types are most likely to have fitness effects.

**Figure 1.**
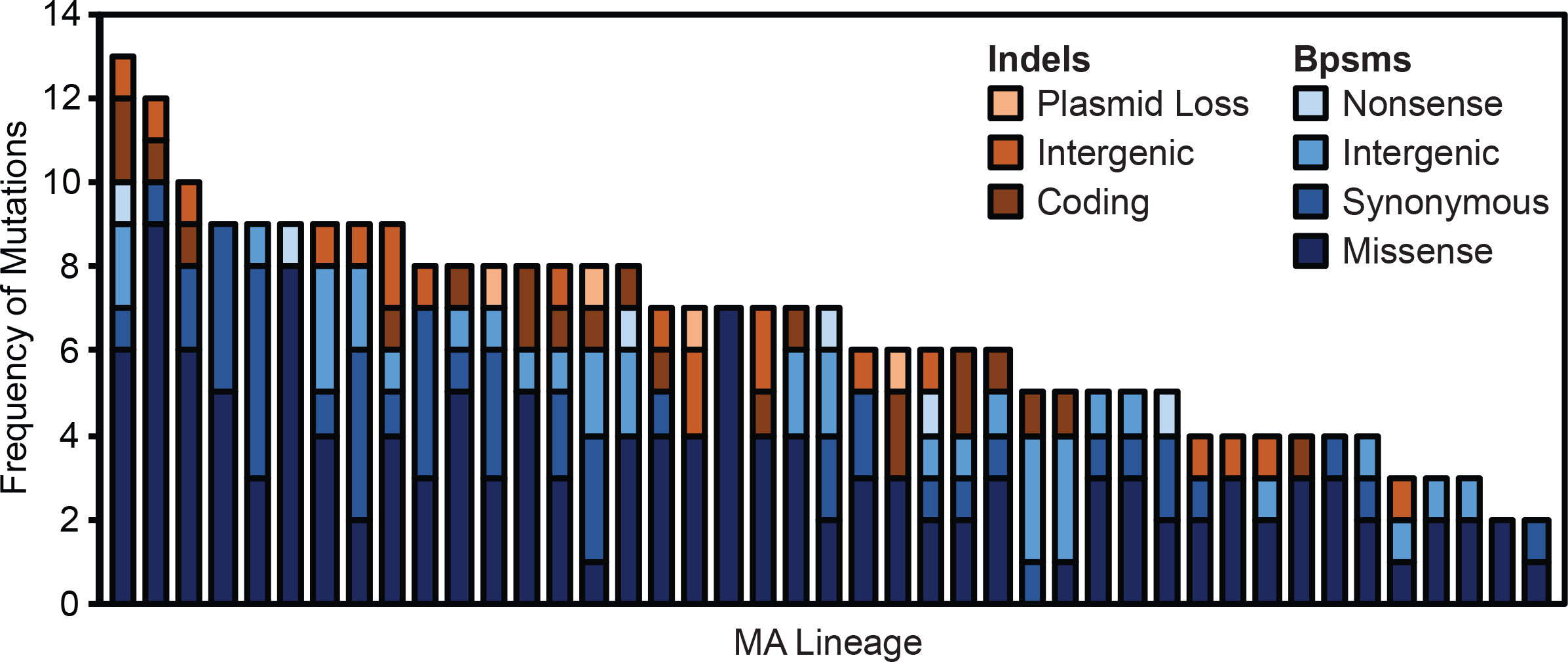
Distribution of base-substitution mutations (bpsms) and insertion-deletion mutations (indels) across the forty-three mutation accumulation lineages derived from *Burkholderia cenocepacia* HI2424 analyzed in this study.Selection Coefficient (s)

**Fitness effects of spontaneous mutations.** Fitness of the final isolate from each lineage was measured by direct competition with the ancestral *B. cenocepacia* HI2424 strain in three different broth culture conditions. TSOY broth is a very permissive medium used to mimic the conditions of the MA experiment, M9MM+CAA is an amino-acid supplemented minimal medium that is less permissive than TSOY but more permissive than a strictly minimal medium, and M9MM is a fully defined minimal medium that is the least permissive of the three environments. In TSOY, 17 lineages had significantly reduced fitness and no lineages had significantly increased fitness (see Figure 2A). The average fitness across all MA lineages in TSOY was −0.024 (0.005) (SEM), with a range of −0.111 to +0.037. Similarly, 13 lineages had significantly reduced fitness in M9MM+CAA and none had significantly increased fitness (see Figure 2B). The average fitness decline and the range across all MA lineages in M9MM+CAA were similar to those observed in TSOY (Average: −0.020 (0.005) (SEM); Range: −0.116 to +0.006). Lastly, we observed 13 lineages with significantly reduced fitness in M9MM, but here, 4 other lineages had significantly increased in fitness (see Figure 2C). Thus, the average fitness decline of the MA lineages in M9MM was only −0.013 (0.005) (SEM), and the range across all lineages was shifted to the right (−.090 to +0.026). Because each lineage harbors multiple mutations, the distributions presented in Figure 2 cannot be used directly to elucidate the distribution of effects of individual spontaneous mutations. However, it is notable that all of the distributions are significantly non-normal (Shapiro Wilk’s Test; TSOY: W = 0.95, p = 0.04; M9MM+CAA: W = 0.68, p = 2.02 • 10^−8^; M9MM: W = 0.73, p = 1.44 • 10^−7^) and the basic properties of the distributions outlined above are similar across environments. Specifically, the majority of lineages have neutral or moderately deleterious fitness and all three distributions have an extended left tail including lineages whose fitnesses have declined more dramatically.

**Figure 2.**
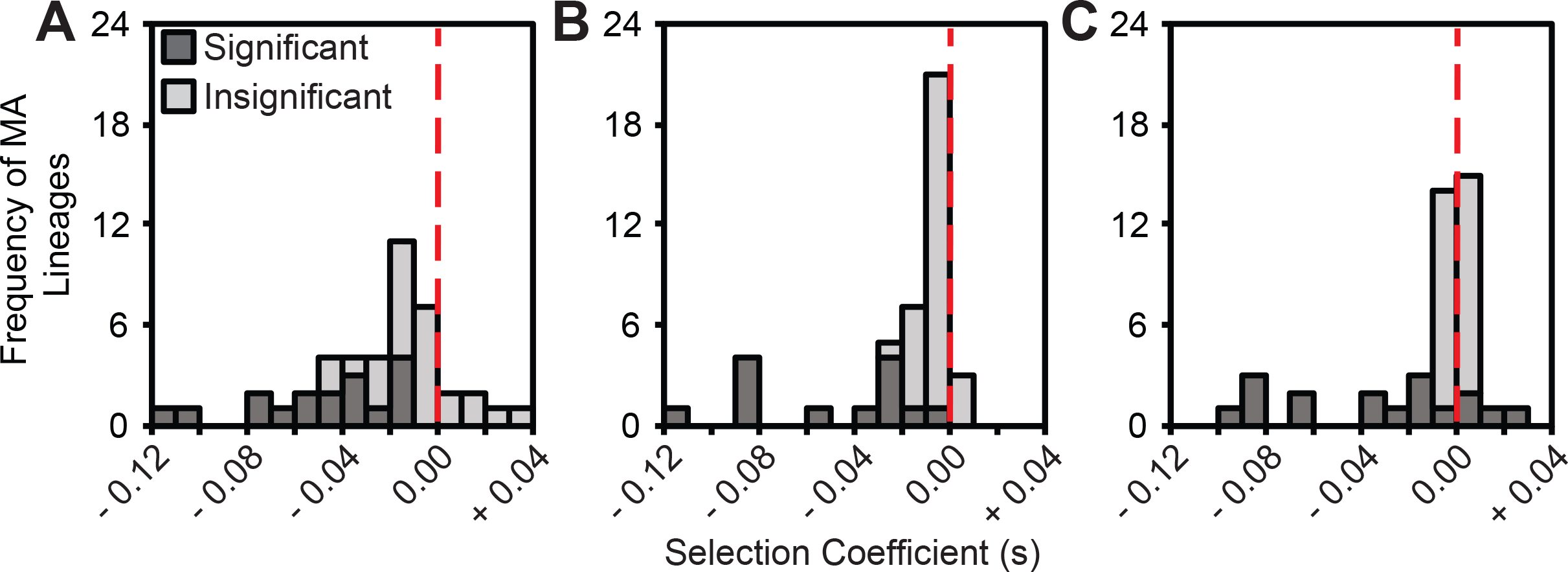
Distribution of the selection coefficients of each *Burkholderia cenocepacia* MA lineage relative to the ancestral *B. cenocepacia* HI2424 strain in tryptic soy broth (A), M9 minimal medium supplemented with casamino acids (B), and M9 minimal medium (C). Significance was determined from independent two-tailed t-tests on four replicate fitness assays for each lineage. P-values were corrected for multiple comparisons using a Benjamini-Hochberg correction, and corrected p-values that remained below 0.05 were considered significant.

Significant positive correlations between the selection coefficients of individual MA lineages across environments suggest that effects of experiment-wide mutational load interacted with these external environments only modestly (see Figure 3). Specifically, linear regressions between the selection coefficients of each lineage in TSOY and both M9MM+CAA and M9MM produced significantly positive relationships (TSOY-M9MM+CAA: F = 17.18, df = 41, p = 0.0002, r^2^ = 0.30; TSOY-M9MM: F = 8.61, df = 41, p = 0.0054, r^2^ = 0.17). Selection coefficients in M9MM+CAA and M9MM were also significantly correlated and explained a greater fraction of the variance than either of the TSOY regressions (F = 124.00, df = 41, p < 0.0001, r^2^ = 0.75), which was expected given that these environments are more similar to each other than either is to TSOY (see Figure 3). However, the fitness of certain MA lineages declined significantly in one environment but not others, suggesting that some mutations in these lineages produced environment-dependent effects. A total of nine lineages were significantly less fit in a single environment (four TSOY, two M9MM+CAA, three M9MM), eight MA lineages were significantly less fit in two of the environments, and six MA lineages were significantly less fit in all three environments. Overall, fitness was significantly influenced by replicate, genotype, environment, and genotype-by-environment interaction (see Table 1). Yet, as noted above, the general properties of the distribution of fitness effects of these MA lineages remained similar across the three tested environments (see Figure 2).

**Table 1.** An experiment-wide ANOVA of the fitness of the 43 final mutation accumulation lines in each of 3 environments, measured in 4 replicates per treatment.

| Effect | df | SS | F | p |
| --- | --- | --- | --- | --- |
| Replicate | 3 | 0.0037 | 10.02 | < 0.0001 |
| Line | 42 | 0.3302 | 63.72 | < 0.0001 |
| Environment | 2 | 0.0190 | 76.90 | < 0.0001 |
| Genotype • Environment | 84 | 0.1166 | 11.25 | < 0.0001 |

**Figure 3.**
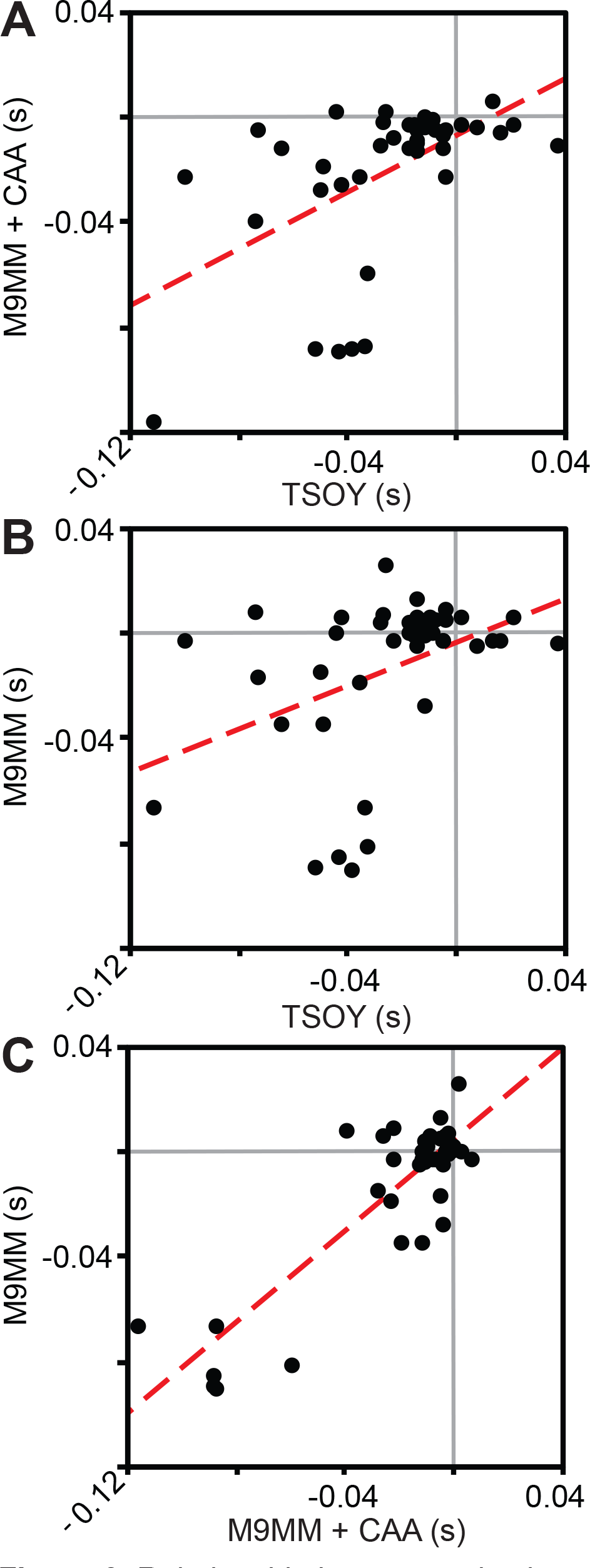
Relationship between selection coefficients of all MA lineages in each of the different pairs of environments. All linear regressions are significant, but much of the variance is unexplained (A: F = 17.18, df = 41, p = 0.0002, r^2^ = 0.2953; B: F = 8.613, df = 41, p = 0.0054, r^2^ = 0.1736; C: F = 124.00, df = 41, p < 0.0001, r^2^ = 0.7515).

Despite acquiring multiple mutations, a number of MA lineages did not have significant fitness differences from the ancestral strain. Further, the number of spontaneous mutations in a line did not correlate with their absolute selection coefficients in any environment (TSOY: F = 1.40, df = 41, p = 0.2434, r^2^ = 0.03; M9MM+CAA: F = 1.35, df = 41, p = 0.2513, r^2^ = 0.03; M9MM: F = 2.96, df = 41, p = 0.0930, r^2^ = 0.07) (see Figure S1). After adding the 11 additional mutations presumed to have been missed in the unanalyzed genomic regions across all lines, we estimate that a total of 290 spontaneous mutations occurred in the experiment, with an average of 6.73 (0.36) (SEM) mutations per lineage. In combination, these results suggest that the fitness effects of a majority of spontaneous mutations were near neutral, or at least undetectable with plate-based laboratory fitness assays. Given the average selection coefficient of each line and the number of mutations that it harbors, we can estimate that the average fitness effect (*s*) of a single mutation was −0.0040 (0.0008) in TSOY, −0.0031 (0.0007) in M9MM+CAA, and −0.0017 (0.0007) (SEM) in M9MM.

Because the fitness of many lineages with multiple mutations did not significantly differ from the ancestor, and because mutation number and fitness were not correlated, this study suggests that most of the significant losses and gains in fitness were caused by rare, single mutations with large fitness effects. Based on this conjecture, we can use the number of lineages that experienced significantly reduced fitness (17 in TSOY, 13 in M9MM+CAA, and 13 in M9MM) divided by the total generations of MA across all lines to estimate the deleterious mutation rate per genome per generation (*U_D_*) in each environment. Estimates of *U_D_* based on this approach are 7.12 × 10^−5^ /genome/generation in TSOY, 5.44 × 10^−5^ /genome/generation in M9MM+CAA, and 5.44 × 10^−5^ /genome/generation in M9MM. The fitness effects of these single deleterious mutations (*s_D_*) in some lineages can also be estimated directly from the fitness of these select lineages relative to the *B. cenocepacia* HI2424 ancestor, and suggest an average *s_D_* of −0.048 (0.007) in TSOY, −0.053 (0.011) in M9MM+CAA, and −0.048 (0.009) in M9MM (SEM). Although we observe no significantly beneficial mutations in TSOY or M9MM+CAA, our data suggest that the beneficial mutation rate (*U_B_*) is 1.68 × 10^−5^ /genome/generation and the average significantly beneficial mutation has a selection coefficient (*s_B_*) of 0.013 (0.005) in M9MM. The primary caveats to these estimates are that they assume that only one of the mutations in each lineage determines its fitness, that none of the lineages that are statistically indistinguishable from the ancestral strain harbor deleterious mutations, and that they do not account for epistasis between the mutations accumulated in each lineage. To attain more precise estimates of *U_D_, s_D_, U_B_*, and *s_B_*, studying spontaneous mutations in different combinations and using more precise fitness assays based on flow-cytometry and/or barcoded sequencing will be especially valuable.

**Genetic basis of deleterious load.** Without sequencing and measuring fitness at intermediate time-points or genetically engineering *B. cenocepacia* HI2424 strains that harbor only single spontaneous mutations, it is difficult to pinpoint which mutations generate the fitness declines in our MA lineages. However, we can examine relationships between the forms of mutational load harbored by each lineage and the fitness of those lineages (Figure 1). The only mutation type that was significantly overrepresented in lineages with reduced fitness in TSOY was the loss of the 0.164-Mb plasmid (χ^2^ = 6.12, d.f. = 1, p = 0.0130). All four lineages that lost the plasmid were significantly less fit in TSOY, with an average selection coefficient of −0.060 (0.007) (SEM). These same four lineages were also less fit in M9MM (χ^2^ = 9.23, d.f. = 1, p = 0.0020; mean s = −0.043 (0.018)), but the deleterious effects of plasmid loss appear to be mitigated in M9MM+CAA, where only one of these lineages had significantly reduced fitness.

Although no other mutation types were significantly overrepresented among lineages that had reduced fitness, we note that there were more coding indels, nonsense bpsms, and missense bpsms than expected in the lineages with reduced fitness in all three environments, with the exception of coding indels in M9MM+CAA (see Table 2). In contrast, intergenic bpsms and indels appear to be evenly distributed between lineages with significantly reduced fitness and those where *s* was not significantly different from 0, suggesting that few if any intergenic mutations from this study have deleterious fitness effects. Similarly, the synonymous bpsms observed in this study do not appear to have deleterious effects, as they are less frequent than expected in lineages with reduced fitness in TSOY and are evenly distributed between neutral and reduced fitness lineages in M9MM+CAA and M9MM (see Table 2). Overall, these results support the expectation that coding indels, nonsense bpsms, and missense bpsms are more likely to have deleterious effects than intergenic and synonymous bpsms.

**Table 2.** Chi-tests comparing the observed number of mutations in lineages with significantly reduced fitness, relative to the expected amounts given the proportion of the total lineages that had significantly reduced fitness.

| Mutation Type | Environment | Observed | Expected | $\chi^2$ | df | p |
| --- | --- | --- | --- | --- | --- | --- |
| Intergenic Bpsm | TSOY | 13 | 14.63 | 0.2996 | 1 | 0.5841 |
| Synonymous Bpsm | TSOY | 16 | 19.37 | 0.9708 | 1 | 0.3245 |
| Missense Bpsm | TSOY | 60 | 55.74 | 0.5374 | 1 | 0.4635 |
| Nonsense Bpsm | TSOY | 4 | 2.37 | 1.8477 | 1 | 0.1741 |
| Intergenic Indel | TSOY | 8 | 7.91 | 0.0018 | 1 | 0.9661 |
| Coding Indel | TSOY | 12 | 8.70 | 2.0736 | 1 | 0.1499 |
| Plasmid Loss | TSOY | 4 | 1.58 | 6.1176 | 1 | 0.0134 |
| Total Mutations | TSOY | 117 | 110.30 | 0.6726 | 1 | 0.4121 |
| Intergenic Bpsm | M9MM+CAA | 12 | 11.19 | 0.0849 | 1 | 0.7708 |
| Synonymous Bpsm | M9MM+CAA | 15 | 14.81 | 0.0033 | 1 | 0.9539 |
| Missense Bpsm | M9MM+CAA | 47 | 42.63 | 0.6427 | 1 | 0.4227 |
| Nonsense Bpsm | M9MM+CAA | 3 | 1.81 | 1.1115 | 1 | 0.2917 |
| Intergenic Indel | M9MM+CAA | 6 | 6.05 | 0.0005 | 1 | 0.9819 |
| Coding Indel | M9MM+CAA | 6 | 6.65 | 0.0914 | 1 | 0.7624 |
| Plasmid Loss | M9MM+CAA | 1 | 1.21 | 0.0519 | 1 | 0.8198 |
| Total Mutations | M9MM+CAA | 90 | 84.35 | 0.5427 | 1 | 0.4613 |
| Intergenic Bpsm | M9MM | 10 | 11.19 | 0.1803 | 1 | 0.6712 |
| Synonymous Bpsm | M9MM | 15 | 14.81 | 0.0033 | 1 | 0.9539 |
| Missense Bpsm | M9MM | 50 | 42.63 | 1.8274 | 1 | 0.1764 |
| Nonsense Bpsm | M9MM | 2 | 1.81 | 0.0274 | 1 | 0.8686 |
| Intergenic Indel | M9MM | 6 | 6.05 | 0.0005 | 1 | 0.9819 |
| Coding Indel | M9MM | 8 | 6.65 | 0.3921 | 1 | 0.5312 |
| Plasmid Loss | M9MM | 4 | 1.21 | 9.2308 | 1 | 0.0024 |
| Total Mutations | M9MM | 95 | 84.35 | 1.9278 | 1 | 0.1650 |

## DISCUSSION

Nearly all prior research on the fitness effects of mutations has studied mutants that have been screened by selection, which presumably purged deleterious variants and enriched beneficial ones. MA experiments are designed to minimize effects of selection to the greatest extent possible, thus capturing most mutations independent of the biases of natural selection. Combining this MA approach with whole genome sequencing and fitness measurements provides the potential to dramatically advance our understanding of the fitness effects of spontaneous mutations in diverse organisms. Here we analyzed the fitness effects of 43 lineages of *B. cenocepacia* that underwent 7 months or >5500 generations of evolution under greatly limited selection. Direct competitions with the common ancestor were conducted in three environments to quantify effects of different forms of mutational load. Despite the duration of MA, many lineages evidently suffered no fitness decline, as they had selection coefficients that did not significantly differ from s = 0. Given that each lineage accumulated between 2-14 mutations, the most likely explanation for these findings is that the vast majority of spontaneous mutations have minimal effects on fitness across this range of environments. Under the assumption that the significant reductions in lineage fitness were driven mostly by single deleterious mutations (Davies *et al.* 1999; Heilbron *et al.* 2014), we also obtain new estimates of the deleterious mutation rate (*U_D_*) and the average effect of deleterious mutations (*s_D_*) in all three environments. These measures reveal that the general features of the distribution of fitness effects are similar between these conditions. The most consistently deleterious mutational event involved loss of the 0.164-Mb plasmid, which reduced fitness in TSOY and M9MM but not M9MM+CAA. Further, nonsense bpsms, missense bpsms, and coding indels were more likely to have contributed to the deleterious mutational load that synonymous bpsms, intergenic bpsms, and intergenic indels.

Although a few select studies have claimed that a substantial fraction of spontaneous mutations are beneficial under certain conditions (Shaw *et al.* 2002; Silander *et al.* 2007; Dickinson 2008), evidence from diverse sources strongly suggests that the effect of most spontaneous mutations is to reduce fitness (Kibota and Lynch 1996; Keightley and Caballero 1997; Fry *et al.* 1999; Vassilieva *et al.* 2000; Wloch *et al.* 2001; Zeyl and de Visser 2001; Keightley and Lynch 2003; Trindade *et al.* 2010; Heilbron *et al.* 2014). Our measurements of selection coefficients in TSOY also suggest that most spontaneous mutations are neutral or deleterious because all lineages whose fitness differs significantly from the ancestor are less fit (*s* = −0.112 to −0.014). The overall distribution of selection coefficients of our lineages in TSOY also has a clear mode near *s* = 0 and among lineages whose selection coefficients cannot be statistically distinguished from s = 0, most are clearly negative (Chi-square test; χ^2^ = 7.54, df = 1, p = 0.0060) (see Figure 2). However, whether this is the only mode in the distribution, or stems from the inability of our MA experiments to fix deleterious mutations with fitness effects below the selection threshold of s = −0.078 remains uncertain.

Whether the environment affects the fitness effects of spontaneous mutations has also been the subject of considerable debate. Specifically, some studies have shown that larger declines in fitness are experienced in harsher environments, while others have not (Martin and Lenormand 2006; Halligan and Keightley 2009; Kraemer *et al.* 2015). In M9MM+CAA and M9MM, we can statistically distinguish fitness effects from s = 0 with greater precision (s < −0.03 in M9MM+CAA and s < −0.01 or s > 0.01 in M9MM) likely because the formulations for these media are more defined. These media are also expected to be more stringent for growth than TSOY because nutrients are more limited. Yet, a similar distribution of fitness effects was observed across our MA lineages in these environments as we observed in TSOY (see Figure 2). Again, most lineages whose selection coefficients are statistically different from 0 in M9MM+CAA and M9MM have reduced fitness, and the only clear mode in the distribution of fitness effects occurs near *s* = 0. However, in M9MM there are four lineages that are significantly more fit than the ancestor, and unlike in TSOY and M9MM+CAA, lineages whose selection coefficients are not significantly different from 0 in M9MM are no more likely to be negative than they are to be positive (Chi-square test; χ^2^ = 0, df = 1, p = 1) (see Figure 2). This suggests that fewer spontaneous mutations are deleterious for fitness in M9MM, possibly because a greater proportion of genes are unused when metabolizing only a single carbon substrate. Furthermore, a number of lineages whose selection coefficients were significantly different from s = 0 in one environment, were near neutral in other environments. Overall, these data suggest that the fitness effects of some individual spontaneous mutations interact with the growth environment, despite the minimal differences in the properties of the distribution of fitness effects among the three environments assayed in this study (see Figure 2; Figure 3).

Quantitative estimates of the rates and fitness effects of spontaneous mutations have received considerable interest because of their fundamental importance for the population genetics of evolving systems (Keightley and Lynch 2003; Eyre-Walker and Keightley 2007; Lynch 2008a; b, 2010; Halligan and Keightley 2009). Specifically, in an effort to unveil some general properties of the distribution of fitness effects of spontaneous mutations, estimates of the average selection coefficient of spontaneous mutations (*s*) (Lynch *et al.* 2008; Halligan and Keightley 2009; Trindade *et al.* 2010; Heilbron *et al.* 2014), the rate of deleterious mutations (*U_D_*) (Kibota and Lynch 1996; Trindade *et al.* 2010), and the average selection coefficient of deleterious mutations (*s_D_*) (Kibota and Lynch 1996; Trindade *et al.* 2010; Heilbron *et al.* 2014) have been derived using both direct and indirect approaches. Here, we estimate that *s* ≅ 0 in all three environments, largely because the vast majority of mutations appear to have near neutral effects on fitness. These estimates are remarkably similar to estimates from studies of MA lines with fully characterized mutational load in *P. aeruginosa* and S. *cereviseae* (Lynch *et al.* 2008; Heilbron *et al.* 2014), but are lower than estimates derived from unsequenced MA lineages (Halligan and Keightley 2009; Trindade *et al.* 2010). However, it is important to consider that while our data suggest that the vast majority of spontaneous mutations in *B. cenocepacia* have very low selection coefficients in the laboratory, it should not imply that all of these mutations are effectively neutral in natural conditions. In fact, sequence analyses in enteric bacteria have revealed that fewer than 2.8% of amino-acid changing mutations are evolving neutrally, and this may be an overestimate due to the presence of adaptive mutations (Charlesworth and Eyre-Walker 2006; Eyre-Walker and Keightley 2007).

From this study, we were also able to estimate the deleterious mutation rates (*U_D_*) and the mean fitness effects of deleterious mutations (*s_D_*) in each environment under the presumption that only one mutation is responsible for the fitness declines experienced by a subset of our lineages. This assumption is warranted given that more than 5000 generations of MA had no significant effects on lineage fitness in the majority of our lineages, and that there was no significant correlation between the number of mutations in a lineage and its fitness. Further, time-series data from prior studies have supported that single, large-effect mutations disproportionately dictate the realized fitness in the majority of MA lineages (Davies *et al.* 1999; Heilbron *et al.* 2014). As was the case with our estimates of *s*, our estimates of *U_D_* and *s_D_* in all three environments are similar to prior estimates in *E. coli* (Kibota and Lynch 1996; Trindade *et al.* 2010) (TSOY: *U_D_* = 7.12 × 10^−5^ /genome/generation, *s_D_* = −0.048; M9MM+CAA: *U_D_* = 5.44 × 10^−5^, *s_D_* = −0.053; M9MM: *U_D_* = 5.44 × 10^−5^, *s_D_* = −0.048). Interestingly, our *U_D_* estimates are all slightly lower than the estimates from previous studies and our *s_D_* estimates are somewhat greater, which is consistent with a failure to differentiate some moderately deleterious mutations from s = 0 in this study. It is likely that both the difficulty of resolving the fitness of mutations of small effects and the effects of selection in our MA experiments on mutations with *s* > 0.078 interfere with our estimates of *U_D_* and *s_D_*.

While the latter selective bias is an inevitable consequence of the MA approach, future studies can enhance resolution of subtle fitness effects by employing flow-cytometry and/or barcoding techniques (Gallet *et al.* 2012; Levy *et al.* 2015; Dillon *et al.* 2016). Furthermore, by extending these methods to a growing database of archived MA-WGS studies in diverse organisms and analyzing the effects of spontaneous mutations in different combinations, we will soon be able to evaluate the extent to which properties of the distribution of fitness effects vary between species and the epistatic deviation associated with mutations in different genetic backgrounds (Elena and Lenski 1997; Silander *et al.* 2007; Dickinson 2008; Schaack *et al.* 2013; Jasmin and Lenormand 2015; Behringer and Hall 2016).

Significantly beneficial mutations were only observed in M9MM and in no lineage did the selective coefficient exceed *s* = +0.03. These limited observations prevent us from performing any detailed analyses on the rate and distribution of effects of beneficial mutations, but they support the notion that beneficial mutations are rare relative to deleterious mutations (Keightley and Lynch 2003). Furthermore, they suggest that most beneficial mutations likely provide only moderate benefits, even though the beneficial mutations that often fix in experimental populations can have large beneficial effects (Lenski *et al.* 1991; Ostrowski *et al.* 2005; Lang *et al.* 2013; Levy *et al.* 2015). As was the case for deleterious mutations, the number and average effects of beneficial mutations observed in this study are likely to be downwardly biased due to the impacts of other mutations in the same MA background that are mostly deleterious, which will further increase the value of sequencing and measuring the fitness of our lineages at intermediate time-points in future studies.

It is a well-established dogma in evolutionary biology that mutations that disrupt coding sequences are most likely to affect fitness, but this has never been quantitatively tested with naturally accumulated mutations. Specifically, mutations that frequently generate non-functional proteins, like nonsense bpsms or coding indels, are expected to have the most deleterious effects, followed by missense bpsms that mostly generate modified proteins, then synonymous and non-coding mutations that do not alter protein sequences. The fitness effects of plasmid gain and loss are less certain, as the size and genetic content of plasmids vary, but they may be energetically expensive to maintain (Smith and Bidochka 1998). Consequently, plasmids may be selectively lost in permissive laboratory environments where maintenance of the plasmid has a fitness cost (Lenski and Bouma 1987; Smith and Bidochka 1998). Our data suggest that although loss of the 0.164-Mb plasmid in *B. cenocepacia* occurs at an appreciable rate in the absence of selection during our MA experiments, it is universally deleterious to lose the plasmid in TSOY and M9MM. However, these effects appear to be mitigated in M9MM+CAA, suggesting that these fitness loses are related to amino acid synthesis. Therefore, these data suggest that in permissive laboratory conditions, the loss of some plasmids can be deleterious and not just advantageous for growth as is widely presumed. Other mutation types were not significantly overrepresented in lineages with significantly reduced fitness (see Table 2), but we do find that there are slightly more nonsense bpsms, missense bpsms, and coding indels than expected in lineages with significantly reduced fitness. This supports expectations that protein-modifying mutations are more likely to affect fitness than synonymous or intergenic mutations, and that most synonymous and intergenic mutations do not measurably affect fitness, even though some synonymous and intergenic mutations can be under selective constraints (Eyre-Walker and Keightley 2007; Bailey *et al.* 2014).

The rate and fitness effects of spontaneous mutations are fundamental quantities that will help explain a number of evolutionary phenomena, including the origin and maintenance of genetic variation in natural populations (Charlesworth *et al.* 1993, 2009; Charlesworth and Charlesworth 1998), the evolution of recombination (Muller 1964; Kondrashov 1988; Otto and Lenormand 2002; Roze and Blanckaert 2014), the evolution of mutator alleles (Sniegowski *et al.* 1997; Tenaillon *et al.* 1999), and the mutational meltdown of small populations (Lande 1994; Lynch *et al.* 1995, 1999; Schwander and Crespi 2009). Although our methods show that the majority of spontaneous mutations in *B. cenocepacia* do not produce detectable fitness effects in any of three laboratory environments, natural selection may be operating on many of these sites in natural populations (Charlesworth and Eyre-Walker 2006). Still, our data suggest that purifying selection may act weakly on much of the variation generated by spontaneous mutation, allowing mutation pressure to substantively influence the genetic variation in natural populations under some conditions. The low frequencies of highly deleterious or beneficial alleles observed in this study may also have implications for our understanding of the evolution of recombination and mutation rates. Specifically, the benefits of reassorting genomes to separate deleterious and beneficial mutations (to avoid Muller’s ratchet), or to combine beneficial mutations and accelerate adaptation (the Fisher-Muller hypothesis) might be reduced under conditions where both deleterious and beneficial mutations are rare (Zeyl and Bell 1997; Otto and Lenormand 2002; de Visser and Elena 2007), like those observed in this study. Mutator alleles may also be tolerated for longer periods if they fail to produce detectable deleterious load, thereby giving them more time to hitchhike to fixation by producing a highly beneficial allele. Indeed, mutator alleles have been frequently observed in experimental (Sniegowski *et al.* 1997), clinical (Mena *et al.* 2008; Oliver 2010; Silva *et al.* 2016), and environmental studies (Hall and Henderson-Begg 2006; Hazen *et al.* 2009). Lastly, the bias for spontaneous mutations to be deleterious rather than beneficial in *B. cenocepacia* is consistent with studies in a number of species (Kibota and Lynch 1996; Keightley and Caballero 1997; Fry *et al.* 1999; Vassilieva *et al.* 2000; Wloch *et al.* 2001; Zeyl and de Visser 2001; Keightley and Lynch 2003; Trindade *et al.* 2010; Heilbron *et al.* 2014), suggesting that gradual mutational meltdown may be inevitable for populations where the efficiency of natural selection is reduced (Sniegowski and Lenski 1995; Lynch *et al.* 1999).

By measuring the fitness effects of MA lineages with fully characterized mutational load, we have performed a uniquely systematic study of the rate and fitness effects of spontaneous mutations with known genetic bases, demonstrating that the vast majority of mutations accumulated in *B. cenocepacia* MA lines are neutral or deleterious for fitness, and that the fitness of individual mutations can be environmentally dependent, even though the general features of the distribution of fitness effects are similar in different environments. Furthermore, we have provided new estimates for several parameters of the distribution of fitness effects of spontaneous mutations derived from genotypes with known mutational load. Although considerable uncertainty remains with respect to the shape and parameters that define the distribution of fitness effects, by extending these methods to a growing database of MA studies, many of which have archived time-series data, we are now poised to dramatically enhance our understanding of the true nature of the distribution of fitness effects of spontaneous mutations.

## ACKNOWLEDGMENTS

This work was supported by the National Science Foundation Career Award (DEB-0845851 to VSC).

## LITERATURE CITED

Agnoli K., Schwager S., Uehlinger S., Vergunst A., Viteri D. F., Nguyen, D. T., Sokol, P. A., Carlier A., Eberl L., 2012 Exposing the third chromosome of Burkholderia cepacia complex strains as a virulence plasmid. Mol. Microbiol. 83: 362–378.

Agrawal A. F., Whitlock, M. C., 2012 Mutation load: The fitness of individuals in populations where deleterious alleles are abundant. Annu. Rev. Ecol. Evol. Syst. 43: 115–135.

Bailey S. F., Hinz A., Kassen R., 2014 Adaptive synonymous mutations in an experimentally evolved Pseudomonas fluorescens population. Nat. Commun. 5: 4076.

Bateman A. J., 1959 The viability of near-normal irradiated chromosomes. Int. J. Radiat. Biol. 1: 170–180.

Baym M., Kryazhimskiy S., Lieberman, T. D., Chung H., Desai, M. M., Kishony R., 2015 Inexpensive multiplexed library preparation for megabase-sized genomes. PLoS One 10: e0128036.

Behringer M. G., Hall, D. W., 2016 The repeatability of genome-wide mutation rate and spectrum estimates. Curr. Genet. 1: 1–6.

Benjamini Y., Hochberg Y., 1995 Controlling the false discovery rate: a practical and powerful approach to multiple testing. J. R. Stat. Soc. 57: 289–300.

Charlesworth B., Morgan, M. T., Charlesworth D., 1993 The effect of deleterious mutations on neutral molecular variation. Genetics 134: 1289–1303.

Charlesworth B., Charlesworth D., 1998 Some evolutionary consequences of deleterious mutations. Genetica 102–103: 3–19.

Charlesworth J., Eyre-Walker A., 2006 The rate of adaptive evolution in enteric bacteria. Mol. Biol. Evol. 23: 1348–1356.

Charlesworth B., Betancourt, A. J., Kaiser, V. B., Gordo I., 2009 Genetic recombination and molecular evolution. In: Cold Spring Harbor Symposia on Quantitative Biology, pp. 177–186.

Chevin L.-M., 2011 On measuring selection in experimental evolution. Biol. Lett. 7: 210–213.

Choi K.-H., Gaynor, J. B., White, K. G., Lopez C., Bosio, C. M., Karkhoff-Schweizer R. R., Schweizer, H. P., 2005 A Tn7-based broad-range bacterial cloning and expression system. Nat. Methods 2: 443–448.

Coenye T., LiPuma, J. J., 2003 Population structure analysis of Burkholderia cepacia genomovar III: varying degrees of genetic recombination characterize major clonal complexes. Microbiology-Sgm 149: 77–88.

Davies E. K., Peters a D., Keightley, P. D., 1999 High frequency of cryptic deleterious mutations in Caenorhabditis elegans. Science 285: 1748–1751.

Dettman J. R., Sztepanacz, J. L., Kassen R., 2016 The properties of spontaneous mutations in the opportunistic pathogen Pseudomonas aeruginosa. BMC Genomics 17: 27–41.

Dickinson W. J., 2008 Synergistic fitness interactions and a high frequency of beneficial changes among mutations accumulated under relaxed selection in Saccharomyces cerevisiae. Genetics 178: 1571–1578.

Dillon M. M., Sung W., Lynch M., Cooper, V. S., 2015 The rate and molecular spectrum of spontaneous mutations in the GC-rich multichromosome genome of Burkholderia cenocepacia. Genetics 200: 935–946.

Dillon M. M., Rouillard, N. P., Dam B. Van, Gallet R., Cooper, V. S., 2016 Diverse phenotypic and genetic responses to short-term selection in evolving Escherichia coli populations. Evolution (N. Y). 70: 586–599.

Elena S. F., Lenski, R. E., 1997 Test of synergistic interactions among deleterious mutations in bacteria. Nature 390: 395–398.

Elena S. F., Ekunwe L., Hajela N., Oden, S. A., Lenski, R. E., 1998 Distribution of fitness effects caused by random insertion mutations in Escherichia coli. Genetica 102: 349–358.

Estes S., Phillips, P. C., Denver, D. R., Thomas, W. K., Lynch M., 2004 Mutation accumulation in populations of varying size: The distribution of mutational effects for fitness correlates in Caenorhabditis elegans. Genetics 166: 1269–1279.

Eyre-Walker A., Keightley, P. D., 2007 The distribution of fitness effects of new mutations. Nat. Rev. Genet. 8: 610–8.

Foster P. L., Lee H., Popodi E., Townes, J. P., Tang H., 2015 Determinants of spontaneous mutation in the bacterium Escherichia coli as revealed by whole-genome sequencing. Proc. Natl. Acad. Sci. 112: E5990–E5999.

Fry J. D., Keightley, P. D., Heinsohn, S. L., Nuzhdin S. V, 1999 New estimates of the rates and effects of mildly deleterious mutation in Drosophila melanogaster. Proc. Natl. Acad. Sci. U. S. A. 96: 574–579.

Gallet R., Cooper, T. F., Elena, S. F., Lenormand T., 2012 Measuring selection coefficients below 10(-3): Method, questions, and prospects. Genetics 190: 175–186.

Hall L. M. C., Henderson-Begg S. K., 2006 Hypermutable bacteria isolated from humans - a critical analysis. Microbiology 152: 2505–2514.

Hall D. W., Mahmoudizad R., Hurd, A. W., Joseph, S. B., 2008 Spontaneous mutations in diploid Saccharomyces cerevisiae: another thousand cell generations. Genet. Res. (Camb). 90: 229–241.

Halligan D. L., Keightley, P. D., 2009 Spontaneous mutation accumulation studies in evolutionary genetics. Annu. Rev. Ecol. Evol. Syst. 40: 151–172.

Hazen T. H., Kennedy, K. D., Chen S., Yi, S. V., Sobecky, P. A., 2009 Inactivation of mismatch repair increases the diversity of Vibrio parahaemolyticus. Environ. Microbiol. 11: 1254–1266.

Heilbron K., Toll-Riera M., Kojadinovic M., Maclean, R. C., 2014 Fitness is strongly influenced by rare mutations of large effect in a microbial mutation accumulation experiment. Genetics 197: 981–990.

Jasmin J., Lenormand T., 2015 Accelerating mutational load is not due to synergistic epistasis or mutator alleles in mutation accumulation lines of yeast. Genetics.

Katju V., Packard, L. B., Bu L., Keightley, P. D., Bergthorsson U., 2015 Fitness decline in spontaneous mutation accumulation lines of Caenorhabditis elegans with varying effective population sizes. Evolution (N. Y). 69: 104–116.

Keightley P. D., 1994 The distribution of mutation effects on viability in Drosophila melanogaster. Genetics 138: 1315–1322.

Keightley P. D., Caballero A., 1997 Genomic mutation rates for lifetime reproductive output and lifespan in Caenorhabditis elegans. Proc. Natl. Acad. Sci. U. S. A. 94: 3823–3827.

Keightley P. D., Lynch M., 2003 Toward a realistic model of mutations affecting fitness. Evolution (N. Y). 57: 683–685.

Kibota T. T., Lynch M., 1996 Estimate of the genomic mutation rate deleterious to overall fitness in E. coli. Nature 381: 694–696.

Kimura M., 1983 The Neutral Theory of Molecular Evolution. Cambridge University Press, Cambridge, New York.

Kondrashov A. S., 1988 Deleterious mutations and the evolution of sexual reproduction. Nature 336: 435–440.

Kraemer S. A., Morgan, A. D., Ness, R. W., Keightley, P. D., Colegrave N., 2015 Fitness effects of new mutations in Chlamydomonas reinhardtii across two stress gradients. J. Evol. Biol. 44: n/a–n/a.

Lande R., 1994 Risk of population extinction from fixation of new deleterious mutations. Evolution (N. Y). 48: 1460–1469.

Lang G. I., Rice, D. P., Hickman, M. J., Sodergren E., Weinstock, G. M., Botstein D., Desai, M. M., 2013 Pervasive genetic hitchhiking and clonal interference in forty evolving yeast populations. Nature 500: 571–574.

Lee H., Popodi E., Tang, H. X., Foster, P. L., 2012 Rate and molecular spectrum of spontaneous mutations in the bacterium Escherichia coli as determined by whole-genome sequencing. Proc. Natl. Acad. Sci. U. S. A. 109: E2774–E2783.

Lenski R. E., Bouma, J. E., 1987 Effects of segregation and selection on instability of plasmid pACYC184 in Escherichia coli B. J. Bacteriol. 169: 5314–5316.

Lenski R. E., Rose, M. R., Simpson, S. C., Tadler, S. C., 1991 Long-term experimental evolution in Escherichia coli .1. Adaptation and divergence during 2,000 generations. Am. Nat. 138: 1315–1341.

Levy S. F., Blundell, J. R., Venkataram S., Petrov, D. A., Fisher, D. S., Sherlock G., 2015 Quantitative evolutionary dynamics using high-resolution lineage tracking. Nature 519: 181–186.

Li H., Durbin R., 2009 Fast and accurate short read alignment with Burrows-Wheeler transform. Bioinformatics 25: 1754–1760.

Li H., Handsaker B., Wysoker A., Fennell T., Ruan J., Homer N., Marth G., Abecasis G., Durbin R., 2009 The sequence alignment/map format and SAMtools. Bioinformatics 25: 2078–2079.

LiPuma J. J., Spilker T., Coenye T., Gonzalez, C. F., 2002 An epidemic Burkholderia cepacia complex strain identified in soil. Lancet 359: 2002–2003.

Long H., Sung W., Miller, S. F., Ackerman, M. S., Doak, T. G., Lynch M., 2014 Mutation rate, spectrum, topology, and context-dependency in the DNA mismatch repair (MMR) deficient Pseudomonas fluorescens ATCC948. Genome Biol. Evol. 7: 262–271.

Long H., Kucukyildirim S., Sung W., Williams E., Lee H., Ackerman M., Doak, T. G., Tang H., Lynch M., 2015 Background mutational features of the radiation-resistant bacterium Deinococcus radiodurans. Mol. Biol. Evol. 32: 2383–2392.

Lynch M., Conery J., Burger R., 1995 Mutation accumulation and the extinction of small populations. Am. Nat. 146: 489–518.

Lynch M., Blanchard J., Houle D., Kibota T., Schultz S., Vassilieva L., Willis J., 1999 Perspective: Spontaneous deleterious mutation. Evolution (N. Y). 53: 645–663.

Lynch M., 2008a Estimation of Nucleotide Diversity, Disequilibrium Coefficients, and Mutation Rates from High-Coverage Genome-Sequencing Projects. Mol. Biol. Evol. 25: 2409–2419.

Lynch M., 2008b The Cellular, Developmental and Population-Genetic Determinants of Mutation-Rate Evolution. Genetics 180: 933–943.

Lynch M., Sung W., Morris K., Coffey N., Landry, C. R., Dopman, E. B., Dickinson, W. J., Okamoto K., Kulkarni S., Hartl, D. L., Thomas, W. K., 2008 A genome-wide view of the spectrum of spontaneous mutations in yeast. Proc. Natl. Acad. Sci. U. S. A. 105: 9272–9277.

Lynch M., 2010 Evolution of the mutation rate. Trends Genet. 26: 345–352.

Mahenthiralingam E., Urban, T. A., Goldberg, J. B., 2005 The multifarious, multireplicon Burkholderia cepacia complex. Nat. Rev. Microbiol. 3: 144–156.

Martin G., Lenormand T., 2006 The fitness effect of mutations across environments: a survey in light of fitness landscape models. Evolution 60: 2413–2427.

Mena A., Smith E. E., Burns, J. L., Speert, D. P., Moskowitz, S. M., Perez, J. L., Oliver A., 2008 Genetic adaptation of Pseudomonas aeruginosa to the airways of cystic fibrosis patients is catalyzed by hypermutation. J. Bacteriol. 190: 7910–7917.

Mukai T., 1964 Genetic Structure of Natural Populations of Drosophila Melanogaster .1. Spontaneous Mutation Rate of Polygenes Controlling Viability. Genetics 50: 1–&.

Muller H. J., 1964 The relation of recombination to mutational advance. Mutat. Res. Mol. Mech. Mutagen. 1: 2–9.

Oliver A., 2010 Mutators in cystic fibrosis chronic lung infection: Prevalence, mechanisms, and consequences for antimicrobial therapy. Int J Med Microbiol 300: 563–572.

Ostrowski E. A., Rozen, D. E., Lenski, R. E., 2005 Pleiotropic effects of beneficial mutations in Escherichia coli. Evolution (N. Y). 59: 2343–2352.

Otto S. P., Lenormand T., 2002 Resolving the paradox of sex and recombination. Nat Rev Genet 3: 252–261.

Perfeito L., Sousa a., Bataillon T., Gordo I., 2014 Rates of fitness decline and rebound suggest pervasive epistasis. Evolution (N. Y). 68: 150–162.

R Development Core Team, 2013 R: A Language and Environment for Statistical Computing.

Roze D., Blanckaert A., 2014 Epistasis, pleiotropy, and the mutation load in sexual and asexual populations. Evolution (N. Y). 68: 137–149.

Schaack S., Allen, D. E., Latta, L. C., Morgan, K. K., Lynch M., 2013 The effect of spontaneous mutations on competitive ability. J. Evol. Biol. 26: 451–6.

Schultz S. T., Lynch M., Willis, J. H., 1999 Spontaneous deleterious mutation in Arabidopsis thaliana. Proc. Natl. Acad. Sci. U. S. A. 96: 11393–8.

Schwander T., Crespi, B. J., 2009 Twigs on the tree of life? Neutral and selective models for integrating macroevolutionary patterns with microevolutionary processes in the analysis of asexuality. Mol. Ecol. 18: 28–42.

Shaw R. G., Byers, D. L., Darmo E., 2000 Spontaneous mutational effects on reproductive traits of Arabidopsis thaliana. Genetics 155: 369–378.

Shaw F. H., Geyer, C. J., Shaw, R. G., 2002 A comprehensive model of mutations affecting fitness and inferences for Arabidopsis thaliana. Evolution 56: 453–463.

Silander O. K., Tenaillon O., Chao L., 2007 Understanding the evolutionary fate of finite populations: the dynamics of mutational effects. PLoS Biol. 5: e94.

Silva I. N., Santos, P. M., Santos, M. R., Zlosnik, J. E. A., Speert, D. P., Buskirk S., Waters C., Cooper, V. S., Moreira, L. M., 2016 Long-term evolution of Burkholderia multivorans during a chronic cystic fibrosis infection reveals shifting forces of selection. mSystems 1: e00029–16.

Smith M. A., Bidochka, M. J., 1998 Bacterial fitness and plasmid loss: the importance of culture conditions and plasmid size. Can. J. Microbiol. 44: 351–355.

Sniegowski P. D., Lenski, R. E., 1995 Mutation and adaptation-the directed mutation controversy in evolutionary perspective. Annu. Rev. Ecol. Syst. 26: 553–578.

Sniegowski P. D., Gerrish, P. J., Lenski, R. E., 1997 Evolution of high mutation rates in experimental populations of E. coli. Nature 387: 703–705.

Sung W., Ackerman, M. S., Miller, S. F., Doak, T. G., Lynch M., 2012 Drift-barrier hypothesis and mutation-rate evolution. Proc. Natl. Acad. Sci. U. S. A. 109: 18488–18492.

Sung W., Ackerman, M. S., Gout J.-F., Miller, S. F., Williams E., Foster P. L., Lynch M., 2015 Asymmetric context-dependent mutation patterns revealed through mutation-accumulation experiments. Mol. Biol. Evol. 32: 1672–1683.

Tenaillon O., Toupance B., Nagard H. Le, Taddei F., Godelle B., 1999 Mutators, population size, adaptive landscape and the adaptation of asexual populations of bacteria. Genetics 152: 485–493.

Traverse C. C., Mayo-Smith L. M., Poltak, S. R., Cooper, V. S., 2013 Tangled bank of experimentally evolved Burkholderia biofilms reflects selection during chronic infections. Proc. Natl. Acad. Sci. U. S. A. 110: E250–E259.

Trindade S., Perfeito L., Gordo I., 2010 Rate and effects of spontaneous mutations that affect fitness in mutator Escherichia coli. Philos. Trans. R. Soc. Lond. B. Biol. Sci. 365: 1177–1186.

Vassilieva L. L., Hook a M., Lynch M., 2000 The fitness effects of spontaneous mutations in Caenorhabditis elegans. Evolution (N. Y). 54: 1234–46.

de Visser J. A. G. M., Elena, S. F., 2007 The evolution of sex: empirical insights into the roles of epistasis and drift. Nat. Rev. Genet. 8: 139–149.

Wloch D. M., Szafraniec K., Borts, R. H., Korona R., 2001 Direct estimate of the mutation rate and the distribution of fitness effects in the yeast Saccharomyces cerevisiae. Genetics 159: 441–452.

Ye K., Schulz, M. H., Long Q., Apweiler R., Ning, Z. M., 2009 Pindel: a pattern growth approach to detect break points of large deletions and medium sized insertions from paired-end short reads. Bioinformatics 25: 2865–2871.

Zeyl C., Bell G., 1997 The advantage of sex in evolving yeast populations. Nature 388: 465–468.

Zeyl C., de Visser, J. A., 2001 Estimates of the rate and distribution of fitness effects of spontaneous mutation in Saccharomyces cerevisiae. Genetics 157: 53–61.

